# An increase in spontaneous activity mediates visual habituation

**DOI:** 10.1101/507814

**Authors:** Jae-eun Kang Miller, Bradley R. Miller, Rafael Yuste

## Abstract

The cerebral cortex is spontaneously active^1,2^, but the function of this ongoing activity remains unclear^3, 4^. One possibility is that spontaneous activity provides contextual information in cortical computations, replaying previously learned patterns of activity^5-8^ that conditions the cortex to respond more efficiently, based on past experience. To test this, we measured the response of neuronal populations in mouse primary visual cortex with chronic two-photon calcium imaging during a visual habituation to a specific oriented stimulus. We unexpectedly found that, during habituation, spontaneous activity increased in neurons across the full range of orientation selectivity, eventually matching that of evoked levels. The increase in spontaneous activity strongly correlated with the degree of habituation. In fact, boosting spontaneous activity with two-photon optogenetic stimulation to the levels of stimulus-evoked activity induced habituation in naïve animals. Our study shows that cortical spontaneous activity is causally linked to habituation, which unfolds by minimizing the difference between spontaneous and stimulus-evoked activity levels, rendering the cortex less responsive. We also show how manipulating spontaneous activity can accelerate this type of learning. We hypothesize that spontaneous activity in visual cortex gates incoming sensory information.

---

To determine learning-related neuronal changes in spontaneous and visually-evoked activity in mouse primary visual cortex (V1), we used a visual habituation paradigm^9^. Mice were presented with 5 trials of 30 seconds of a pre-stimulus period with a gray screen followed by 100 seconds of drifting gratings with a single orientation daily for 7 consecutive days (Fig. 1a, b). This protocol induces visual habituation, as reflected by a decrease in movement in response to the stimulus^9^. In a control group, the same drifting gratings were presented on day 1 and day 7, but from day 2 to day 6 drifting gratings with different orientations were presented in a random order each day. We measured running speed as a behavioral output (Fig. 1c) and then calculated habituation index from it (1-normalized running speed during 5 seconds after the onset of a visual stimulus, see the Methods).

**Fig. 1:**
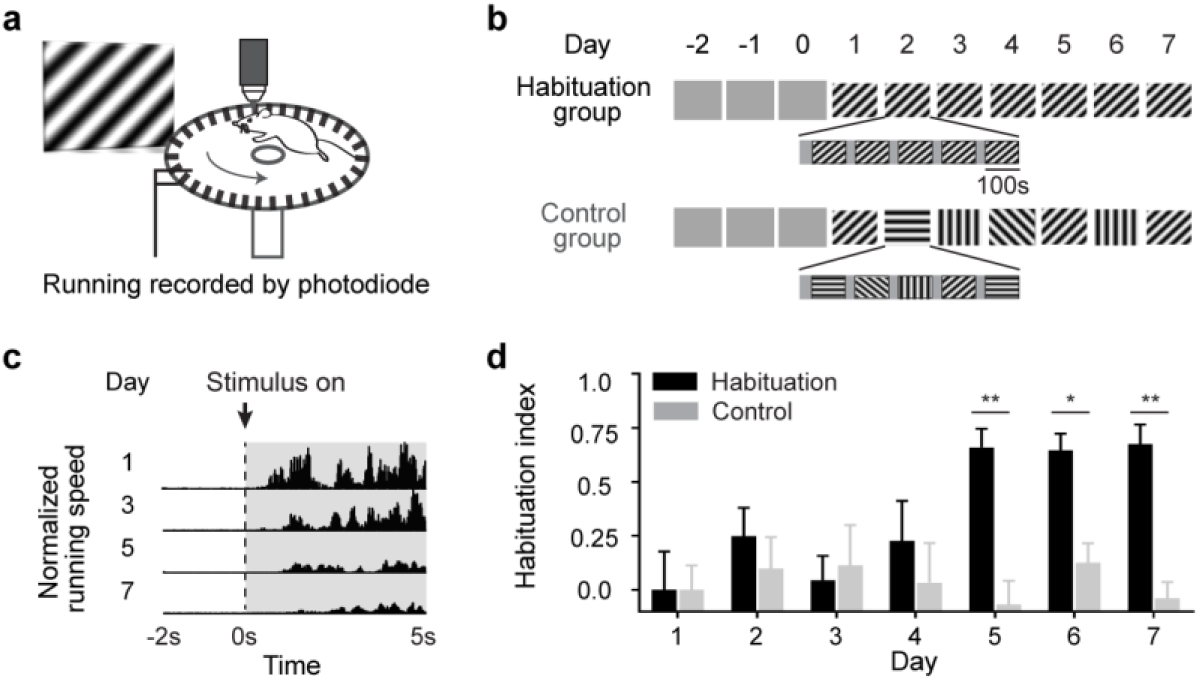
Behavioral habituation to the presentation of drifting oriented gratings. **a**, Illustration of a head-fixed awake chronic two-photon imaging setup. Mice were presented with visual stimulation with drifting gratings and their running speed was recorded by a photodiode. Head fixation was omitted from the drawing for clarity. **b**, Illustration of visual stimulation protocol. Mice became accustomed to head fixation for 3 days while being presented with a gray screen. Over the next 7 days, mice were presented with 5 trials of drifting gratings with a single orientation daily for 10 min. One orientation was randomly selected for each mouse. In a control group, drifting gratings with the same orientation as the habituation group were presented on day 1 and day 7, but from day 2 to day 6 drifting gratings with different orientations were presented in a random order each day. **c**, Example of normalized running speed during 2 seconds prior to and 5 seconds after the onset of visual stimulation from day 1 through day 7. Visual stimulation lasted for 100 seconds for each trial. **d**, Habituation index was calculated from normalized running speed (1-normalized running speed during 5 seconds after the onset of a visual stimulus, see the Methods for the details). n = 14 mice for habituation group; n = 11 mice for control group; *p < 0.05 and **p < 0.005 by two-way ANOVA with Sidak’s multiple comparisons test. Data are presented as mean + s.e.m.

We found that mice habituated to a familiar visual stimulus after 5 days, and the control group did not (Fig. 1d). Habituation was specific to a visual stimulus and not to the behavioral setup, since the control group that were exposed to the same behavioral setup did not habituate. Also, the increase in habituation index in the habituation group was not due to an overall change in locomotion because normalized running speed during the 100 seconds of visual stimulation did not change (Extended Data Fig. 1a). These results demonstrate that our behavioral paradigm can be used to measure the neuronal changes underlying visual habituation.

We hypothesized that habituation will reshape spontaneous cortical activity to become more similar to visually evoked activity, as a form of visual plasticity. To test this hypothesis, we imaged the activity of the same neurons in layer 2/3 of V1 during visual habituation on day 1 and day 7 (Fig. 2a). On each imaging day, spontaneous pre-trial activity was also measured for 10 minutes while mice were kept under total darkness prior to visual stimulation. Interestingly, we observed that neuronal activity during the pre-stimulus periods seemed to be noticeably increased on day 7. To confirm this, we analyzed population activity during the pre-stimulus and visual stimulus periods (Fig. 2a). We found that activity during the pre-stimulus period increased significantly after habituation (34.7% increase on day 7 compared to day 1) whereas the evoked activity during visual stimulus did not change (Fig. 2b). As a result, the difference in the population activity between pre-stimulus and visual stimulus periods on day 1 disappeared on day 7 after habituation. Remarkably, the spontaneous population activity during the pre-trial period also increased after habituation (40.4% increase on day 7 compared to day 1; Fig. 2c).

**Fig. 2:**
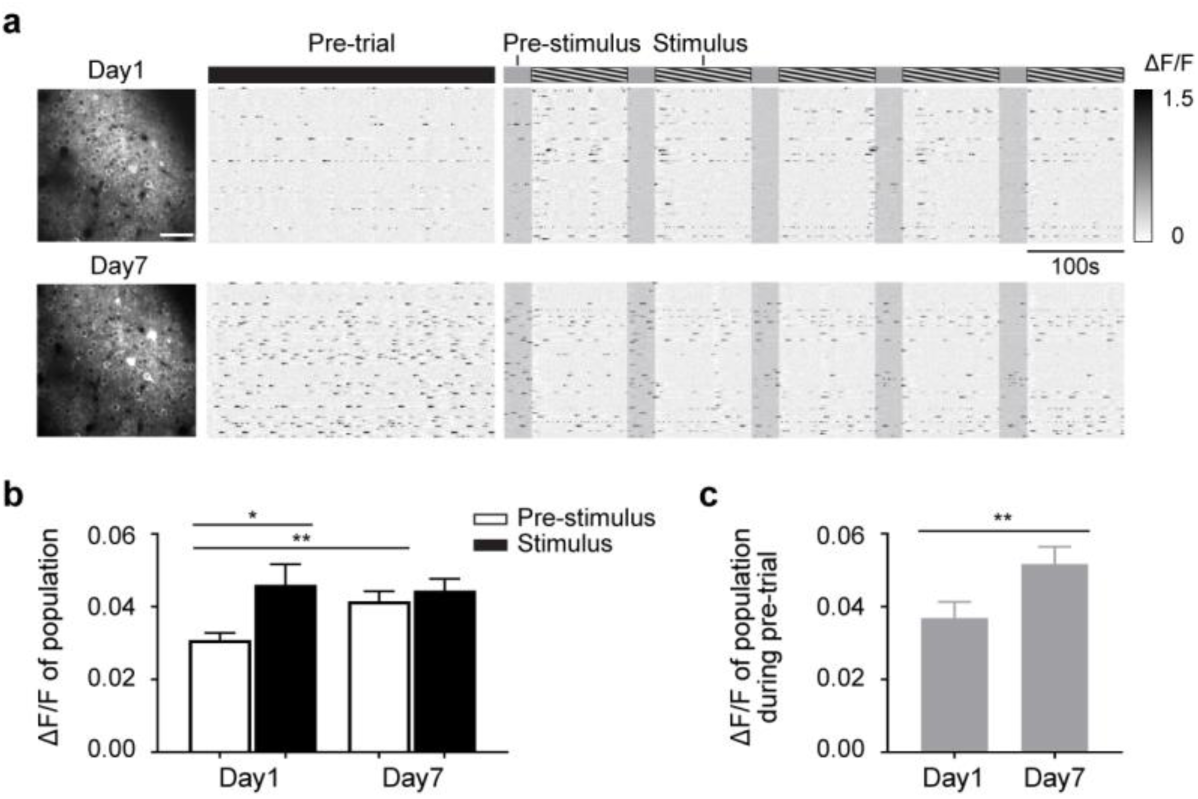
Habituation increases spontaneous activity. **a**, Left, an example of averaged calcium imaging frame of cells in layer 2/3 on day 1 and day 7 in an awake mouse. Right, raster plots of the neuronal activity from the same 120 neurons during pre-trial, pre-stimulus, and visual stimulus periods on day 1 and day 7. **b**, Mean ΔF/F of population during pre-stimulus and visual stimulus periods on day 1 and day 7. n = 7 mice; *p < 0.05 by one-way ANOVA with Tukey’s multiple comparisons test. **c**, Mean ΔF/F of population during pre-trial period on day 1 and day 7. n = 7 mice; **p < 0.01 by paired t test. Data are presented as mean + s.e.m.

The increases in spontaneous population activity during pre-stimulus and pre-trial periods were specific to mice that habituated because population activity did not change on day 7 in the control group (Extended Data Fig. 2). Also, the increases in population activity during pre-stimulus and pre-trial periods were not due to the overall changes in animals’ locomotion because the mean running speed did not change during pre-stimulus or pre-trial period (Extended Data Fig. 1b,c). These results demonstrate that habituation specifically increases cortical spontaneous activity.

This habituation-induced increase in pre-stimulus activity could derive from neurons that are selective for the stimulus since they were stimulated preferentially. This would suggest that stimulus-evoked neuronal ensembles are being primed in preparation for the expected stimulus^10^. To test this, we measured the orientation selectivity of the 23±02% of imaged cells which significantly increased neuronal activity during pre-stimulus period on day 7 (Extended Data Fig. 3a,b, blue). Surprisingly, we found that they are were mixed population, spanning the full range of orientation selectivity (Fig. 3a). The distribution of preferred orientations of cells with increased activity during the pre-stimulus period was indistinguishable from that of all other cells (Fig. 3b). The same was true for cells that increase activity during the pre-trial period after habituation (Extended Data Fig. 3c, red). Thus, it appears that spontaneous activity in V1 increases nonspecifically after habituation.

**Fig. 3:**
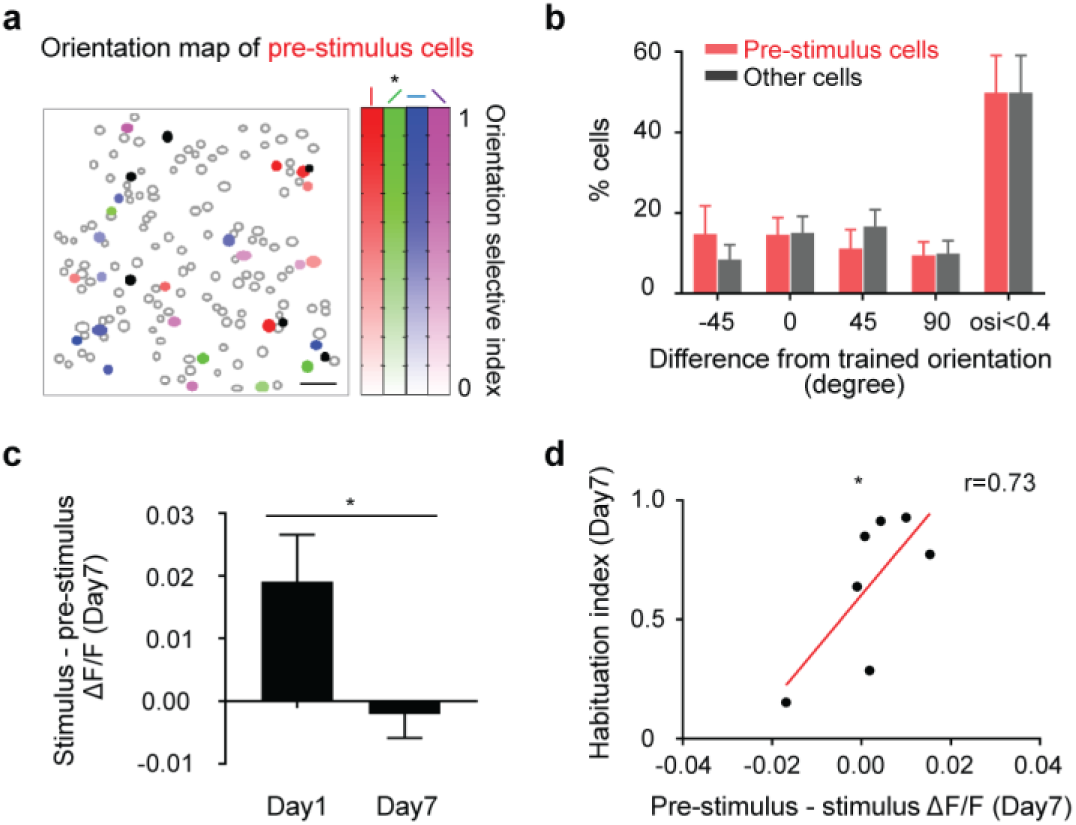
Habituation correlates with decrease in difference between pre-stimulus and stimulus-induced activity. **a**, Example of 150 cells imaged on day 1 and 7. Filled circles indicate the cells that significantly increased ΔF/F during pre-stimulus period on day 7, compared to day 1. These cells were defined as pre-stimulus cells. Orientation index of pre-stimulus cells is color-coded. Black filled circles indicate pre-stimulus cells with no preferred orientation. Star indicates the orientation of a trained visual stimulus. **b**, Distributions of preferred orientations of pre-stimulus cells and the other cells. n = 7 mice; p = 0.36 by two-way ANOVA with Sidak’s multiple comparisons test. **c**, Mean population ΔF/F during stimulus period subtracted by mean population ΔF/F during pre-stimulus period on day 1 and day 7. n = 7 mice; *p < 0.05 by paired t test. **d**, Mean population ΔF/F during pre-stimulus period subtracted by mean population ΔF/F during stimulus period on day 7 significantly correlated with habituation index on day 7. r = 0.73; n = 7 mice; *p < 0.05 by Pearson’s correlation coefficient. Data are presented as mean + s.e.m.

Since habituation increased population activity during the pre-stimulus period, but not during visual stimulus period, the difference between pre-stimulus activity and stimulus-induced neuronal activity significantly decreased after habituation (Fig. 3c). Because of this, we hypothesized that the equalization of pre-stimulus activity to stimulus-evoked activity may drive habituation. Consistent with this hypothesis, we found a strong correlation between the equalization of pre-stimulus activity to stimulus-evoked activity and habituation index (Fig. 3d).

We then set out to examine if there is a causal relationship between the equalization of pre-stimulus activity to stimulus-evoked activity and habituation. To test this, we mimicked the increase in pre-trial and pre-stimulus spontaneous activity using two-photon optogenetics^11^ and examined the effect of this manipulation on behavior (Fig. 4a,b). We simultaneously imaged V1 activity during optogenetic stimulation^12^ to confirm that we were indeed mimicking the natural habituation-induced change in spontaneous activity. Mice were injected with GCaMP6f and C1V1 in V1 seven or eight weeks prior to the optogenetic stimulation experiment. On the experimental day, baseline activity was measured for 10 min prior to performing optogenetic stimulation. Mice then underwent the same habituation paradigm described in Fig. 1 with the addition of optogenetic stimulation during the pre-stimulus and pre-trial periods. Control mice underwent the same experimental procedure except that they were injected only with GCaMP6f. The distribution of orientation selectivity of optogenetically stimulated neurons was indistinguishable from that of neurons that increase spontaneous activity during habituation (Extended Data Fig. 4).

**Fig. 4:**
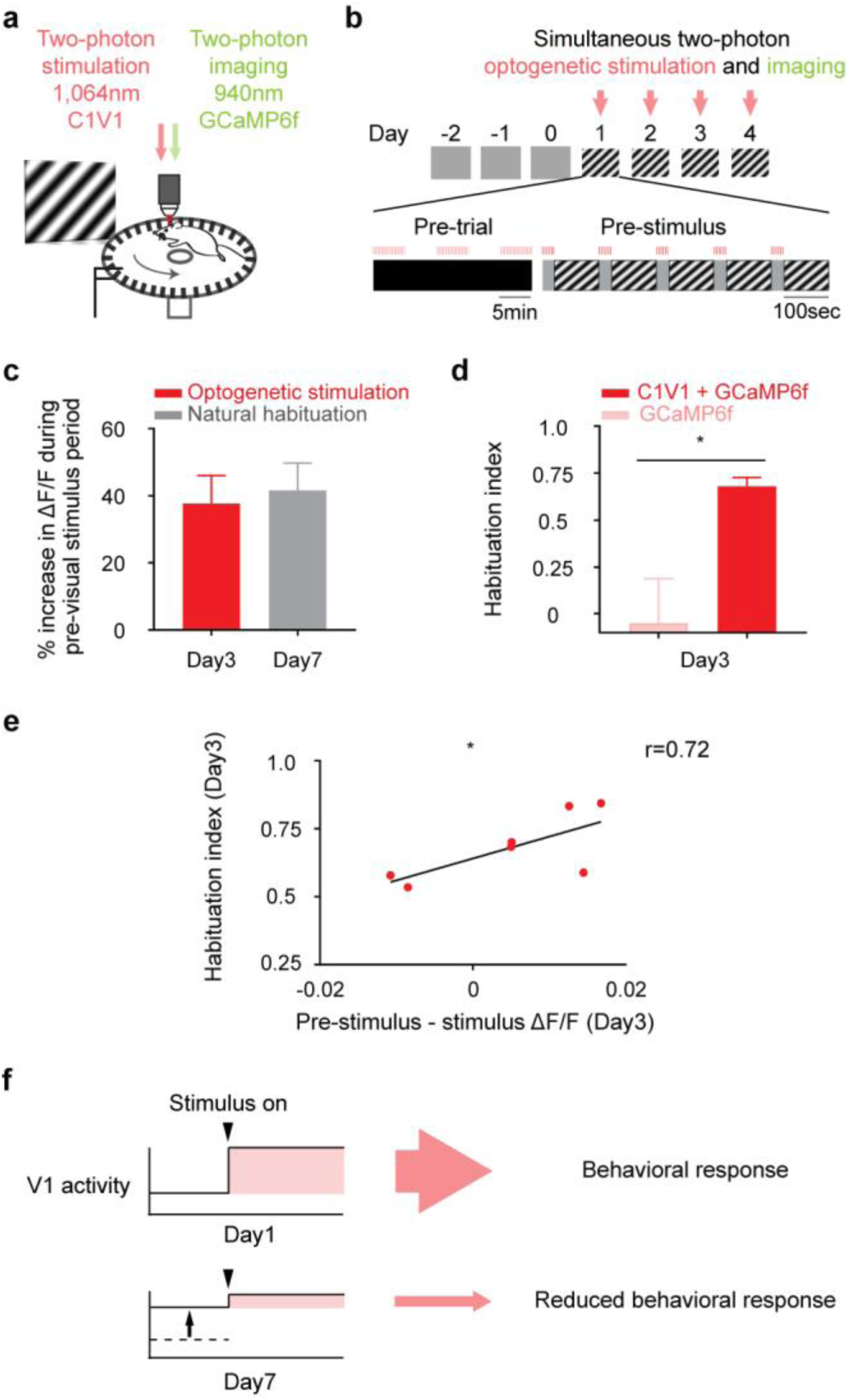
Increasing spontaneous activity accelerates habituation. **a**, Illustration of a head-fixed setup with simultaneous two-photon imaging and two-photon optogenetic stimulation. **b**, Mice became accustomed to head fixation for 3 days while being presented with gray screen. Over the next 4 days, mice were presented with 5 trials of drifting gratings with a single orientation daily as described in Fig. 1. In addition, two-photon optogenetic stimulation was performed during pre-trial and pre-stimulus periods. Calcium imaging was performed simultaneously. **c**, Mean % increase in population ΔF/F during pre-visual stimulus period on day 3 and day 7, compared to population ΔF/F on day 1. A gray bar represent data from mice that underwent natural habituation and a red bar represents data from mice that underwent optogenetic stimulation during pre-trial and pre-stimulus period in addition to a visual stimulus. n = 7 mice for both groups; p = 0.74 by unpaired t test. **d**, Mice habituated to a familiar visual stimulus on day 3. Light red bars represent data from the control group in which mice were injected with GCaMP6f only and dark red bar represent data from the experimental group in which mice were injected with GCaMP6f and C1V1. n = 7 mice for both groups; *p < 0.05 by unpaired t test. **e**, Mean population ΔF/F during pre-visual stimulus period subtracted by mean population ΔF/F during visual stimulus period on day 3 was significantly correlated with habituation index on day 3 from the experimental group in which mice were underwent optogenetic stimulation during pre-trial and pre-stimulus period in addition to a visual stimulus. r = 0.72; n = 7 mice; *p < 0.05 by Pearson’s correlation coefficient. **f**, A proposed model illustrating a circuit basis of visual habituation. When an animal is exposed to a visual stimulus for the first time, stimulus-induced neuronal activity leads to behavioral response. After repeated exposure to the visual stimulus, spontaneous activity increases (a black arrow). As a result, stimulus-induced neuronal activity relative to pre-visual stimulus activity is reduced and this results in a reduced behavioral response. By combining the repeated exposure to a visual stimulus with optogenetic stimulation in V1, we could accelerate visual habituation. Data are presented as mean + s.e.m.

On the first day of optogenetic stimulation, pre-visual stimulus activity levels reached that of day 3 of natural habituation (Extended Data Fig. 5a). Importantly, habituation does not yet occur on day 3 of natural habituation (see Fig. 1d). By day 3 of optogenetic stimulation, we were able to boost pre-visual stimulus V1 activity to the level seen on day 7 of natural habituation (Fig. 4c). In this way, we were able to compare the impact of increasing activity to a level seen before habituation (day 1 of optogenetic stimulation, day 3 of habituation activity level) to a level seen after habituation (day 3 of optogenetic stimulation, day 7 of habituation activity level).

We found that optogenetically increasing pre-visual stimulus V1 activity to that seen after natural habituation induced habituation. On day 3 of optogenetic stimulation, mice reached the full 7 day level of habituation (Fig. 4d). Control mice displayed no habituation (Fig. 4d). Mice did not display habituation on day 1 of optogenetic stimulation, when pre-visual stimulus activity only reached that seen on day 3 of natural habituation (Extended Data Fig. 5b). To probe this further, we took advantage of small inter-animal differences in optogenetic activation and habituation in the optogenetic cohort. We found a strong correlation between degree of increase in activity with optogenetics and the degree of habituation (Fig. 4e). While habituation accelerated in all animals, animals with greater stimulated activity, relative to visually evoked activity, displayed greater habituation (Fig. 4e). Thus, we found a “dose response” relationship of optogenetically increasing pre-visual stimulus activity and habituation. Taken together, this supports the model that the increase in pre-visual stimulus V1 activity to visual stimulus levels drives habituation.

Here we report that cortical spontaneous activity is causally related to habituation. With two-photon calcium imaging, we monitored the activity of the V1 neurons as visual habituation unfolded and unexpectedly discovered a sizable increase in spontaneous V1 activity. We found no such increase in visually evoked activity. Furthermore, spontaneous activity only increased after the repetition of a familiar stimulus. It did not increase after a sequence of novel stimuli. We also found that spontaneous activity increases to the level of stimulus-evoked activity. This equalization is a signature of habituation– it’s strongly correlated with the degree of habituation. We hypothesized that the equalization of spontaneous and stimulus-evoked activity could be causally linked to habituation. We directly tested this with two-photon optogenetics and found that boosting pre-visual stimulus activity levels to that of visual stimulus levels induced habituation. Thus, this increase in spontaneous activity to evoked levels may represent a previously unknown mechanism of learning (Fig, 4f).

This modulation in spontaneous activity is a novel form of cortical plasticity. The increase in spontaneous activity that we found after habituation differs from changes in activity that occur after instrumental learning, where learned sequences of activity replay at rest and may form a kind of internal practice^8, 13, 14^. In contrast, after habituation, we see a widespread increase in V1 activity across the entire spectrum of orientation selectivity. Remarkably, this increase in spontaneous activity is present as soon as the mouse is placed in the habituated context, even prior to the first visual stimulus of the session.

Our findings raise two questions for future study. First, what is the mechanism underlying the increase in spontaneous activity? This could result from disinhibition to increase excitatory drive. A leading candidate is the top down disinhibition by the cingulate cortex. In mouse, the cingulate cortex projects directly to V1 and this input has a net inhibitory effect in V1 activity^15^. Therefore, a decreased input from the cingulate cortex can boost pyramidal neuron activity in V1 via disinhibition. Furthermore, the cingulate is a key regulator of visual attention^16-20^, and thereby a decrease in attention could increase spontaneous activity in V1. Second, how could an increase in spontaneous activity result in habituation? We propose that prior to habituation, a rapid increase in V1 activity from baseline after the presentation of a novel stimulus is a signal for behavioral activation. This rapid increase could be read into and acted on by a motor program^21^. Alternatively, it could be relayed to a regulator of arousal, such as the locus coeruleus^22, 23^, and increase locomotion through an increase in arousal. The learning-induced increase in spontaneous activity would cancel out this rapid increase in V1 activity and results in behavioral habituation.

Regardless of the exact upstream and downstream mechanisms, our results demonstrate one novel function for ongoing spontaneous activity, as gating sensory inputs. Spontaneous activity is found in all cortical areas, and is also a property of all nervous systems, even in cnidarians^24^. Finally, our findings provide a direct demonstration that manipulations of spontaneous activity can accelerate a form of learning, a result that could have clinical and educational implications.

## Methods

### Animals, Surgery, and Training

All experimental procedures were carried out in accordance with Columbia University institutional animal care guidelines. Experiments were performed on C57BL/6 mice at the age of postnatal day (P) 70–P150. Twenty two mice were used for the behavior experiment (14 mice for the habituation group and 8 mice for the control group) and 13 out of 22 mice were imaged using chronic calcium imaging (7 mice for the habituation group and 6 mice for the control group). Fourteen mice were used for the two-photon photostimulation experiment (7 mice for the experimental group and 7 mice for the control group).

Virus injection, head plate implantation, and craniotomy were carried out in that order over the course of 4 to 8 weeks. In the first surgery (virus injection), 3- 7 weeks prior to the first imaging session, mice were anesthetized with isoflurane (initially 3% (partial pressure in air) and reduced to 1%–2%). Carprofen (5 mg/kg) was administered intraperitoneally. A small window was made through the skull above left V1 using a dental drill (coordinates from lambda: X = −250, Y = 30 μm) taking care not to pierce the dura mater. A glass capillary pulled to a sharp micropipette was advanced with the stereotaxic instrument. For chronic calcium imaging only, 400nl solution of 3:1 diluted AAV1.Syn.GCaMP6f.WPRE.SV40 (obtained from the University of Pennsylvania Vector Core) was injected into putative layer 2/3 at a rate of 80 nl/min at a depth of 250 μm from the pial surface using a UMP3 microsyringe pump (World Precision Instruments). For simultaneous two-photon optogenetics and calcium imaging, 650nl solution of mixed AAV1-syn-GCaMP6f and AAVDJ-CaMKII-C1V1-(E162T)-TS-p2A-mCherry-WPRE (1.5:5 ratio; obtained from Stanford Neuroscience Gene Vector and Virus Core) was injected using the same procedure. Our lab previously reported that 40-60% of the cells co-expressed both viruses using this injection method^25^. After surgery animals received carprofen injections for 2 days as post-operative pain medication.

Approximately 3 weeks (for chronic calcium imaging only) or 7 weeks (for simultaneous two-photon optogenetics and calcium imaging) after virus injection, mice were anesthetized as previously described and a titanium head plate was attached to the skull centered on the virus injection site using dental cement. Mice were allowed to recover for a day in their home cage and given carprofen intraperitoneally (5mg/kg) for 2 days. Mice then underwent training to maneuver with their head fixed approximately 1 inch above a rotating wheel while being presented with gray screen for 30 min daily for 3 days. After the first training session, mice could distribute weight on the wheel evenly and appeared calm (grooming, locomotion). This head-fixed awake preparation allows mice to move freely.

After being accustomed to head-fixation for 3 days, mice were anesthetized again with isoflurane. Dexamethasone sodium phosphate (2 mg/kg) and enrofloxacin (4.47 mg/kg) were administered subcutaneously. Carprofen (5 mg/kg) was administered intraperitoneally. A small circle of the skull centered on the virus injection site (approximately 2 mm in diameter) was removed using a dental drill and the dura mater was carefully removed using a fine forceps. A circular glass coverslip (3 mm in diameter) was put on top of the exposed brain and secured using Krazy glue. After surgery animals received carprofen injections for 2 days as post-operative pain medication. Mice were then allowed to recover for a day in their home cage before the first imaging session.

### Visual Habituation

Visual stimuli were generated using the MATLAB Psychophysics Toolbox^26^ and displayed on a liquid crystal display monitor (19-inch diameter, 60-Hz refresh rate) positioned 15 cm from the right eye, roughly at 45° to the long axis of the animal. Pre-trial spontaneous calcium signals were measured for 10 min in the dark prior to the visual stimulation session (pre-trial period). The imaging setup was completely enclosed with blackout fabric (Thorlabs). After pre-trial spontaneous calcium signals were collected, mice were presented with 5 trials of mean luminescence gray screen for 30 seconds (pre-stimulus period) followed by full-field sine wave gratings at a single orientation drifting in one direction (100% contrast, 0.05 cycles per degree, and two cycles per second) for 100 seconds (visual stimulus period). Orientation was randomly chosen for each mouse. Visual stimulation was repeated for 7 days. In a control group, drifting gratings at a single orientation were presented on day 1 and day 7, but from day 2 to day 6 drifting gratings with different orientations were presented in a random order each day. To calculate tuning curves of imaged cells, mice were presented with an additional visual stimulation session at the end of a visual habituation experiment on day 1 and day 7. Full-field sine wave grating stimuli (100% contrast, 0.05 cycles per degree, two cycles per second) drifting in eight different directions were presented in random order for 5 seconds, followed by 5 seconds of mean luminescence gray screen (10 repetitions).

Previously, a behavioral output of a visual habituation was reliably measured in mice as reduced fidget behavior in response to the onset of a visual stimulus in a head-fixed restraint condition^9^. Similarly, a reduced running speed at the onset of a visual stimulus was used as a behavioral output in our head-fixed freely moving condition. Running was tracked using a photodiode coupled to a custom made circuit board to provide a voltage output. Small pieces of black tape (0.5 cm width) were placed under a rotating wheel with 0.5cm intervals and a photodiode was positioned in a rotating wheel as shown in Fig. 1a. As mice pass a piece of black tape, it generates a shift in a voltage signal and a number of shifts in a unit of time provides a running speed.

To calculate a stimulus-induced running speed we analyzed voltage signals from 2 seconds before stimulus onset until 5 seconds after stimulus onset^9^. Voltage signals were first rectified and a mean speed during 5 second after stimulus onset was calculated. Next, a mean speed was normalized to a baseline mean speed during 2 seconds prior to stimulus onset to determine a stimulus-induced running speed and then averaged over 5 trials. To account for the running difference among mice, a stimulus-induced running speed was normalized by the mean of mice and then normalized to the first day to emphasize running changes relative to the first day. Finally, habituation index was calculated as 1-normalized running speed (a maximum habituation index is 1).

### Two-Photon Calcium Imaging and Two-Photon Optogenetic Stimulation

Activity of cortical neurons was recorded by imaging changes of fluorescence with a two-photon microscope (Bruker; Billerica, MA) and a Ti:Sapphire laser (Chameleon Ultra II; Coherent) at 940 nm through a 25x objective (water immersion, N.A. 1.05, Olympus). Resonant galvanometer scanning and image acquisition (frame rate 30.206 or 3.246 fps, 512 × 512 or 256 × 256 pixels, ∼250μm beneath the pial surface) were controlled by Prairie View Imaging software.

Simultaneous two-photon imaging and optogenetic stimulation were performed with two different femtosecond-pulsed lasers attached to a commercial microscope. An imaging laser (λ = 940 nm) was used to excite a genetically encoded calcium indicator (GCaMP6f) while an optogenetic stimulation laser (λ = 1064 nm) was used to excite a red shifted opsin (C1V1) that preferentially responds to longer wavelengths. The two laser beams on the sample are individually controlled by two independent sets of galvanometric scanning mirrors.

Population optogenetic stimulation was performed by synchronizing the photostimulation galvanometers with the imaging galvanometers integrated into the dual beam microscope, i.e. the optogenetic stimulation beam was scanned across the whole field of view. This generates a 4Hz stimulation in each pixel. We decided to use the laser power of 5.7mW and the stimulation duration of 500ms for the rest of optogenetic stimulation experiments based on our previous studies^25, 27^.

In the optogenetic stimulation experiment, the baseline pre-trial activity was measured for 10 min prior to performing photostimulation. After measuring the baseline activity, neurons expressing C1V1 were stimulated during 15 minutes of pre-trial period and during the entire pre-stimulus period. During the pre-trial period, the photostimulation beam was scanned across the whole field of view with the duration of 500 ms, followed by 10 sec interval for 28 trials in one session lasting for 5 min. Three sessions were repeated with 5 min interval between sessions. During the pre-stimulus period, the photostimulation beam was scanned across the whole field of view with the duration of 500 ms, followed by 4 sec interval for 6 trials lasting for the entire pre-stimulus period.

### Image Analysis

Image analysis was performed using ImageJ and built-in and custom-built software in MATLAB. Images were first converted to TIFF format and registered to correct for *x–y* motion using Turboreg plug-in in ImageJ^28^. Initial image processing was carried out using custom-written software in MATLAB (Caltracer 2.5, available at our laboratory website). Regions of interest (ROIs) were drawn around each cell using a semi-automated algorithm based on fluorescence intensity (mean projection), florescence change (standard-deviation projection), and cell size and shape and were adjusted by visual inspection. All pixels within each ROI were averaged to give a single time course, and ΔF/F was calculated by subtracting each value with the mean of the lower 50% of previous 10-s values and dividing it by the mean of the lower 50% of previous 10-s values. Neurons with noisy signal with no apparent calcium transient were detected by visual inspection and excluded from further analysis. Neuropil contamination was removed by first selecting a spherical neuropil shell (6.2-μm thickness) surrounding each neuron and then subtracting the average signal of all pixels within the spherical neuropil shell, excluding adjacent ROIs and pixels within 0.3 μm surrounding ROIs, from the average signal of all pixels within the ROI.

To analyze the orientation selective index (OSI), we used data recorded during an additional imaging session on day 1 in which mice were presented with full-field sine wave grating stimuli (100% contrast, 0.05 cycles per degree, two cycles per second) drifting in eight different directions in random order for 5 seconds, followed by 5 seconds of mean luminescence gray screen (10 repetitions). Average ΔF/F was taken as the response to each grating stimulus. Responses from 10 trials were averaged to obtain an orientation-tuning curve or matrix. The preferred orientation was taken as the modulus of the preferred direction to 180°. The OSI was calculated as (R_best_ – R_ortho_)/(R_best_ + R_ortho_), where R_best_ is the best direction and R_ortho_ is the average of responses to the direction orthogonal to the best direction^29^. Cells with an OSI <0.4 were considered to be unselective for orientation.

### Statistical Analysis

Statistical tests were performed in Prism (Graphpad) software as described in the figure legends.

## Acknowledgments

Laboratory members for comments, help and virus injections. Stanford Neuroscience Gene Vector and Virus Core for AAVdj virus. Supported by the National Eye Institute (DP1EY024503, R01EY011787, K99EY024653). This material is based upon work supported by, or in part by, the U. S. Army Research Laboratory and the U. S. Army Research Office (Contract W911NF-12-1-0594, MURI). The authors declare no competing financial interests. J.K.M., B.R.M. and R.Y. conceptualized this work. J.K.M. and B.R.M. analyzed the data. J.K.M. performed the experiments. J.K.M., B.R.M. and R.Y. wrote the original draft. J.K.M., B.R.M. and R.Y. reviewed and edited the paper. R.Y. directed the study and secured funding. All data are archived at the NeuroTechnology Center at Columbia University.

## Extended Data Figures

**Extended Data Fig. 1.**
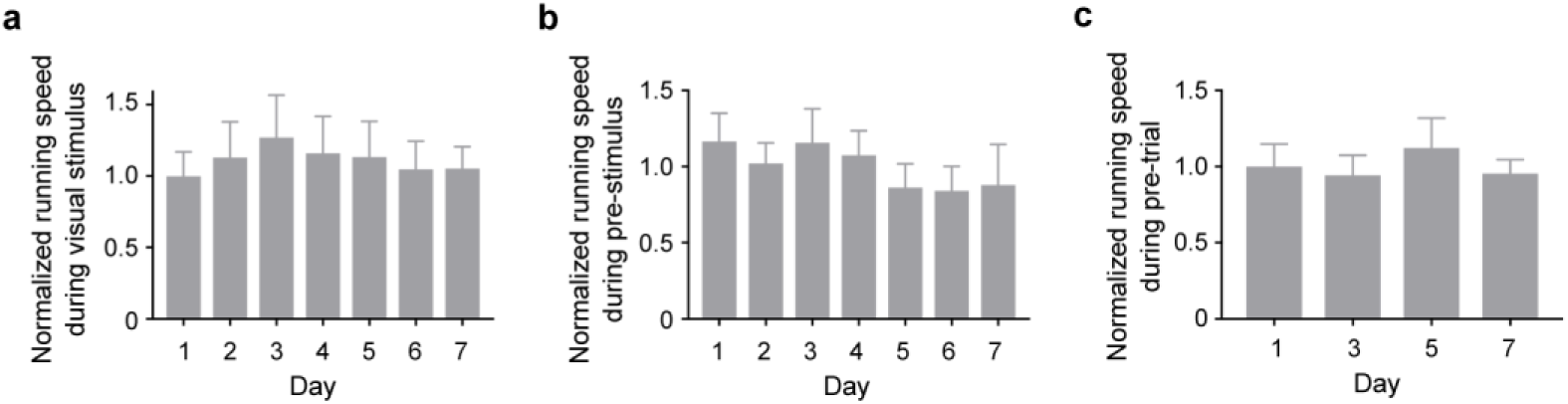
Overall running speed did not change during visual stimulus, pre-stimulus, and pre-trial periods from day 1 through day 7. Normalized running speed (**a**) during 100 seconds of the entire visual stimulus (**b**) during pre-stimulus and (**c**) during pre-trial periods from day 1 to day 7. n = 14 mice; p = 0.70, p = 0.96, and p = 0.75, respectively, by one-way ANOVA with Tukey’s multiple comparisons test. Data are presented as mean + s.e.m.

**Extended Data Fig. 2.**
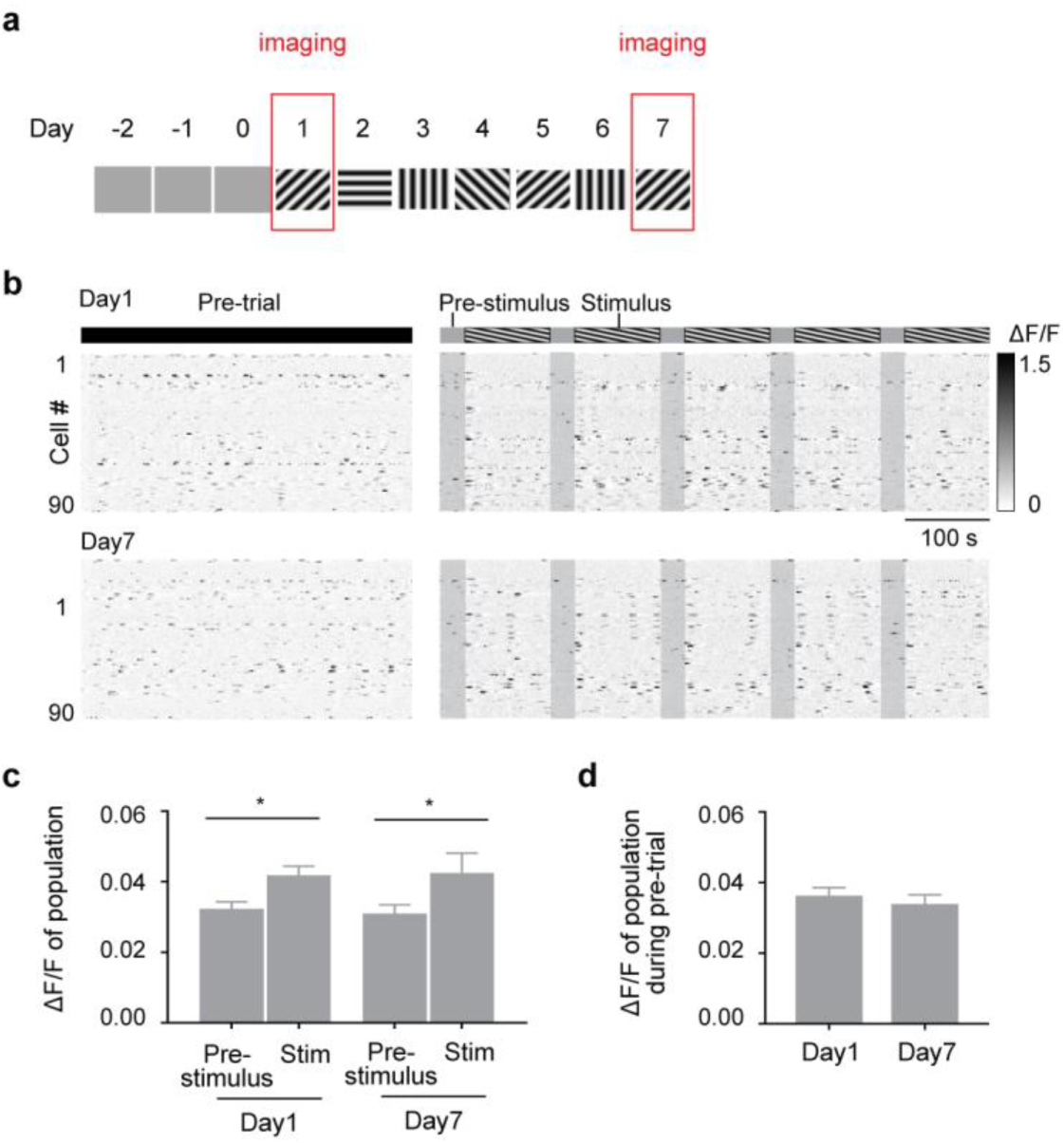
Population activity during pre-trial, pre-stimulus, and visual stimulus periods does not change in a control group. **a**, Illustration of visual stimulation protocol in a control group. Mice became accustomed to head fixation for 3 days while being presented with gray screen and drifting gratings with a same orientation was presented on day 1 and day 7, but from day 2 to day 6 drifting gratings with different orientations were presented in a random order each day as described in Fig. 1. **b**, Raster plots of the neuronal activity from the same 90 neurons in an awake mouse during pre-trial, pre-stimulus, and visual stimulus periods on day 1, top, and day 7, bottom. **c**, Mean population ΔF/F during pre-stimulus and visual stimulus periods on day 1 and day 7. n = 6 mice; *p < 0.05 by one-way ANOVA with Tukey’s multiple comparisons test. **d**, Mean population ΔF/F during pre-trial period on day 1 and day 7. n = 6 mice; p = 0.49 by paired t test. Data are presented as mean + s.e.m.

**Extended Data Fig. 3.**
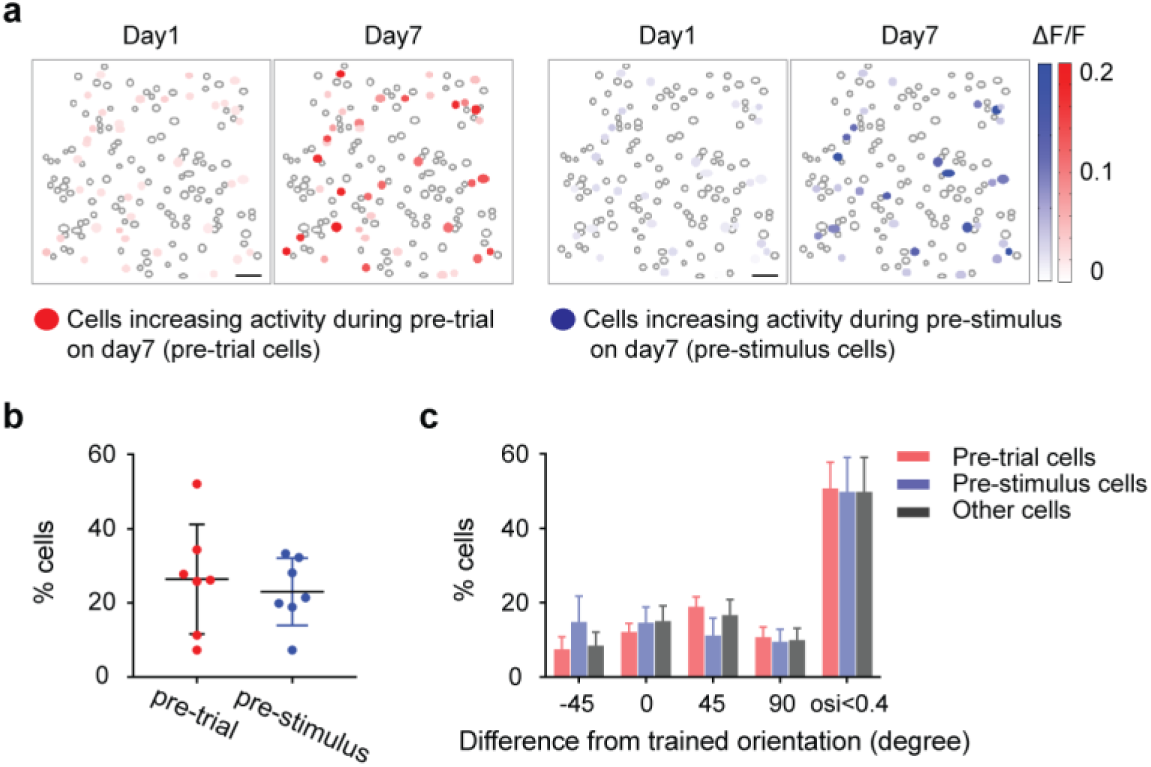
Neurons across the full range of orientation selectivity increase spontaneous activity after habituation. **a**, Example of 150 cells imaged on day 1 and day 7. Red and blue filled circles indicate the cells that significantly increased ΔF/F (color-coded) during pre-trial and pre-stimulus periods, respectively, on day 7, compared to day 1. These cells were defined as pre-trial and pre-stimulus cells, respectively. **b**, Percentage of cells that belong to pre-trial and pre-stimulus cells. Data are presented as mean ± s.d. **c**, Distributions of preferred orientations of pre-trial, pre-stimulus, and the other cells. n = 7 mice; p = 0.91 by two-way ANOVA with Sidak’s multiple comparisons test. Data are presented as mean + s.e.m.

**Extended Data Fig. 4.**
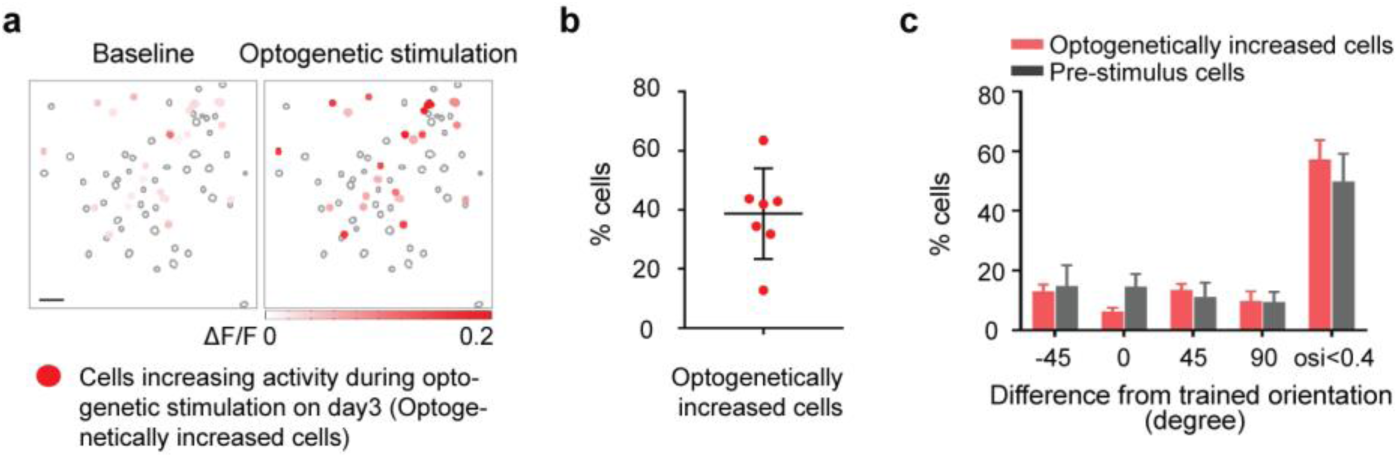
Two-photon optogenetic stimulation mimicking the habituation-induced change in spontaneous activity. **a**, Example of 68 cells imaged. Red filled circles indicate the cells that significantly increased ΔF/F (color-coded) during optogenetic stimulation on day 3, compared to the baseline prior to optogenetic stimulation on day 1. These cells were defined as optogenetically increased cells. **b**, Percentage of cells that belong to optogenetically increased cells. Data are presented as mean ± s.d. **c**, Distributions of preferred orientations of optogenetically increased cells and pre-stimulus cells that were shown in Fig. 2. n = 7 mice; p = 0.13 by two-way ANOVA with Sidak’s multiple comparisons test. Data are presented as mean + s.e.m.

**Extended Data Fig. 5.**
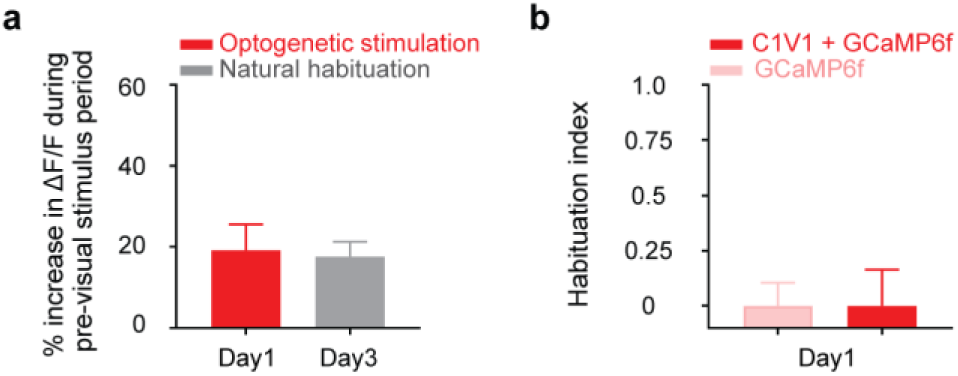
Boosting pre-visual stimulus V1 activity to the level seen prior to habituation (day 3) did not induce habituation. **a**, Mean % increase in population ΔF/F during pre-visual stimulus period on day 1 and day 3, compared to baseline population ΔF/F on day 1. A gray bar represent data from mice that underwent natural habituation and a red bar represents data from mice that underwent optogenetic stimulation during pre-trial and pre-stimulus period in addition to a visual stimulus. n = 7 mice for both groups; p = 0.83 by unpaired t test. **b**, Mice did not habituate on day 1 of optogenetic stimulation. Light red bars represent data from the control group in which mice were injected with GCaMP6f only and dark red bar represent data from the experimental group in which mice were injected with GCaMP6f and C1V1. n = 7 mice for both groups; p = 0.99 by unpaired t test. Data are presented as mean + s.e.m.

## References

1 Berger, H. Über das Elektrenkephalogramm des Menschen. Archiv für Psychiatrie und Nervenkrankheiten 87, 527–570. (1929).

2 Fox, M. D. & Raichle, M. E. Spontaneous fluctuations in brain activity observed with functional magnetic resonance imaging. Nat Rev Neurosci 8, 700–711, doi:nrn2201 [pii] 10.1038/nrn2201 (2007).

3 Raichle, M. E. A paradigm shift in functional brain imaging. The Journal of neuroscience: the official journal of the Society for Neuroscience 29, 12729– 12734, doi:10.1523/JNEUROSCI.4366-09.2009 (2009).

4 Harmelech, T. & Malach, R. Neurocognitive biases and the patterns of spontaneous correlations in the human cortex. Trends in cognitive sciences 17, 606–615, doi:10.1016/j.tics.2013.09.014 (2013).

5 Kenet, T., Bibitchkov, D., Tsodyks, M., Grinvald, A. & Arieli, A. Spontaneously emerging cortical representations of visual attributes. Nature. 425, 954–956 (2003).

6 Miller, J. E., Ayzenshtat, I., Carrillo-Reid, L. & Yuste, R. Visual stimuli recruit intrinsically generated cortical ensembles. Proceedings of the National Academy of Sciences of the United States of America 111, E4053–4061, doi:10.1073/pnas.1406077111 (2014).

7 Berkes, P., Orban, G., Lengyel, M. & Fiser, J. Spontaneous cortical activity reveals hallmarks of an optimal internal model of the environment. Science 331, 83–87, doi:10.1126/science.1195870 (2011).

8 Ji, D. & Wilson, M. A. Coordinated memory replay in the visual cortex and hippocampus during sleep. Nature neuroscience 10, 100–107, doi:10.1038/nn1825 (2007).

9 Cooke, S. F., Komorowski, R. W., Kaplan, E. S., Gavornik, J. P. & Bear, M. F. Visual recognition memory, manifested as long-term habituation, requires synaptic plasticity in V1. Nature neuroscience 18, 262–271, doi:10.1038/nn.3920 (2015).

10 SanMiguel, I., Widmann, A., Bendixen, A., Trujillo-Barreto, N. & Schroger, E. Hearing silences: human auditory processing relies on preactivation of sound-specific brain activity patterns. The Journal of neuroscience: the official journal of the Society for Neuroscience 33, 8633–8639, doi:10.1523/JNEUROSCI.5821-12.2013 (2013).

11 Packer, A. M. et al. Two-photon optogenetics of dendritic spines and neural circuits. Nature methods, doi:10.1038/nmeth.2249 (2012).

12 Yang, W., Carrillo-Reid, L., Bando, Y., Peterka, D. S. & Yuste, R. Simultaneous two-photon imaging and two-photon optogenetics of cortical circuits in three dimensions. Elife 7, doi:10.7554/eLife.32671 (2018).

13 Eagleman, S. L. & Dragoi, V. Image sequence reactivation in awake V4 networks. Proceedings of the National Academy of Sciences of the United States of America 109, 19450–19455, doi:10.1073/pnas.1212059109 (2012).

14 Guidotti, R., Del Gratta, C., Baldassarre, A., Romani, G. L. & Corbetta, M. Visual Learning Induces Changes in Resting-State fMRI Multivariate Pattern of Information. The Journal of neuroscience: the official journal of the Society for Neuroscience 35, 9786–9798, doi:10.1523/JNEUROSCI.3920-14.2015 (2015).

15 Zhang, S. et al. Selective attention. Long-range and local circuits for top-down modulation of visual cortex processing. Science 345, 660–665, doi:10.1126/science.1254126 (2014).

16 Kastner, S. & Ungerleider, L. G. Mechanisms of visual attention in the human cortex. Annual review of neuroscience 23, 315–341, doi:10.1146/annurev.neuro.23.1.315 (2000).

17 Corbetta, M. & Shulman, G. L. Control of goal-directed and stimulus-driven attention in the brain. Nature reviews. Neuroscience 3, 201–215, doi:10.1038/nrn755 (2002).

18 Posner, M. I. & Petersen, S. E. The attention system of the human brain. Annual review of neuroscience 13, 25–42, doi:10.1146/annurev.ne.13.030190.000325 (1990).

19 Wu, D. et al. Persistent Neuronal Activity in Anterior Cingulate Cortex Correlates with Sustained Attention in Rats Regardless of Sensory Modality. Scientific reports 7, 43101, doi:10.1038/srep43101 (2017).

20 Kim, J., Wasserman, E. A., Castro, L. & Freeman, J. H. Anterior cingulate cortex inactivation impairs rodent visual selective attention and prospective memory. Behavioral neuroscience 130, 75–90, doi:10.1037/bne0000117 (2016).

21 Goodale, M. A. Transforming vision into action. Vision research 51, 1567– 1587, doi:10.1016/j.visres.2010.07.027 (2011).

22 Hagan, J. J. et al. Orexin A activates locus coeruleus cell firing and increases arousal in the rat. Proceedings of the National Academy of Sciences of the United States of America 96, 10911–10916 (1999).

23 Sara, S. J. & Bouret, S. Orienting and reorienting: the locus coeruleus mediates cognition through arousal. Neuron 76, 130–141, doi:10.1016/j.neuron.2012.09.011 (2012).

24 Dupre, C. & Yuste, R. Non-overlapping Neural Networks in Hydra vulgaris. Curr Biol 27, 1085–1097, doi:10.1016/j.cub.2017.02.049 (2017).

25 Carrillo-Reid, L., Yang, W., Bando, Y., Peterka, D. S. & Yuste, R. Imprinting and recalling cortical ensembles. Science 353, 691–694, doi:10.1126/science.aaf7560 (2016).

26 Brainard, D. H. The Psychophysics Toolbox. Spatial vision 10, 433–436 (1997).

27 Yang, W., Carrillo-Reid, L., Bando, Y., Peterka, D. S. & Yuste, R. Simultaneous two-photon optogenetics and imaging of cortical circuits in three dimensions. eLife 7, doi:10.7554/eLife.32671 (2018).

28 Thevenaz, P., Ruttimann, U. E. & Unser, M. A pyramid approach to subpixel registration based on intensity. IEEE transactions on image processing: a publication of the IEEE Signal Processing Society 7, 27–41, doi:10.1109/83.650848 (1998).

29 Hofer, S. B. et al. Differential connectivity and response dynamics of excitatory and inhibitory neurons in visual cortex. Nature neuroscience 14, 1045–1052, doi:10.1038/nn.2876 (2011).

